# Chronic ethanol ingestion impairs *Drosophila melanogaster* health in a microbiome-dependent manner

**DOI:** 10.1101/217240

**Authors:** James Angus Chandler, Lina Victoria Innocent, Isaac L. Huang, Jane L. Yang, Michael B. Eisen, William B. Ludington

## Abstract

Ethanol is one of the worlds most abused drugs yet the impacts of chronic ethanol consumption are debated. Ethanol is a prevalent component in the diets of diverse animals and can act as a nutritional source, behavior modulator, and a toxin. The source of ethanol is microbes, which can both produce and degrade ethanol, and the gut microbiome has been associated with differential health outcomes in chronic alcoholism. To disentangle the various and potentially interacting roles of bacteria and ethanol on host health, we developed a model for chronic ethanol ingestion in the adult fruit fly, *Drosophila melanogaster*, which naturally consumes a diet between 0 and 5% ethanol. We took advantage of the tractability of the fly microbiome, which can be experimentally removed to separate the direct and indirect effects of commensal microbes. We found that moderate to heavy ethanol ingestion decreased lifespan and reproduction, without causing inebriation. These effects were more pronounced in flies lacking a microbiome, but could not be explained by simple bacterial degradation of ethanol. However, moderate ethanol ingestion increased reproduction in bacterially-colonized flies, relative to bacteria-free flies. Ethanol decreased intestinal stem cell turnover in bacterially-colonized flies and decreased intestinal barrier failure and increased fat content in all flies, regardless of microbiome status. Analysis of host gene expression finds that ethanol triggers the innate immune response, but only in flies colonized with bacteria. Taken together we show that, chronic ethanol ingestion negatively impacts fly health in a microbiome-dependent manner.

## Introduction

Ethanol is common in the diets of many animals and is also among the most abused drugs in the world. Naturally fermenting diets can contain appreciable amounts of ethanol, which is consumed by a variety of animals including primates, birds, bats, treeshrews, and insects (Hockings et al. 2015; Mazeh et al. 2008; Wiens et al. 2008; Sánchez et al. 2004). The common fruit fly, *Drosophila melanogaster*, naturally consumes ethanol and has long been used as a model for investigating the effects of ethanol on animals (Devineni & Heberlein 2013). Flies have an attraction to ethanol (Devineni & Heberlein 2009; Ja et al. 2007) and display many hallmarks of human alcoholism including tolerance, addiction, and withdrawal (Kaun et al. 2011; Devineni & Heberlein 2009; Ghezzi et al. 2014; Robinson et al. 2012). While the developmental effects of ethanol have been studied in fly larvae (McClure et al. 2011; Logan-Garbisch et al. 2015, the role of long-term oral ingestion of moderate ethanol in adult flies has not been investigated as previous studies focused on larval development and adult intoxication through ethanol vapor. To fill this gap, we developed a *Drosophila* model of chronic ethanol ingestion.

*Drosophila* is a powerful model system for investigating the commensal animal microbiome (Broderick & Lemaitre 2012; Douglas 2018). Flies can be cleared of their microbial communities so that both direct and indirect effects of the microbiome can be investigated (Koyle et al. 2016). Experiments investigating the role of the microbiome can be done on a large scale in flies, testing many variables in parallel (Wong et al. 2014; Gould et al. 2018). Studies have shown that commensal bacteria affect many components of fly fitness and physiology and many of these effects are seen only in a diet-specific context (Shin et al. 2011; Storelli et al. 2011).

Ethanol can be both produced and consumed by microbes, and therefore play a key role in host exposure. In humans, the microbiome is implicated in many of ethanol’s negative consequences, although the relative roles of direct ethanol-induced damage and indirect damage through ethanol’s ability to change the microbiota composition are unclear (Chen & Schnabl 2014; Hartmann et al. 2015). Here we use flies deconstruct the complex interplay between host, microbiome, and ethanol. We find that fly fitness in the presence of ethanol is heavily influenced by the microbiome, and that many aspects of fly biology, including intestinal homeostasis, lipid content, and the immune response, are mediated by microbes following ethanol ingestion. Taken together, our newly developed model of chronic ethanol ingestion in flies provides insight into how the animal microbiome modulates the effects of dietary ethanol.

## Methods

### Fly stocks, husbandry, and creation of ethanol media

All experiments used *Wolbachia*‒free *D. melanogaster* Canton-S strain (Bloomington Line 64349) as previously described (Obadia et al. 2017). Flies were maintained at 25°C with 60% humidity and 12-hour light/dark cycles on autoclaved glucose-yeast medium (10% glucose, 5% Red Star brand active dry yeast, 1.2% agar, 0.42% proprionic acid). Flies were three to six days old before bacterial or ethanol treatments were applied (i.e. all flies were bacteria-free and raised on 0% ethanol diets at birth). Bacteria-free flies were generated by sterilizing dechorionated embryos (Ridley et al. 2013). Bacteria-free stocks were kept for several generations and checked regularly for presence of yeasts, bacteria, and known viruses. Bacterially-colonized flies were created by allowing approximately 50 normally-colonized young adults (from unmanipulated lab stocks) to seed autoclaved media with their frass for about 10 minutes, removing these flies, and then introducing bacteria-free flies. 0% to 15% ethanol media was made by adding 100% ethanol to autoclaved glucose-yeast medium after it had cooled to 50°C. Vials were stored under equivalent ethanol vapor pressure to reduce evaporation until use. Because we were interested in the toxic, rather than nutritional, effects of ethanol, and because the caloric value of ethanol is not easily comparable to that of sugars (Xu et al. 2012), we did not adjust amount of glucose in an attempt to create an isocaloric diet (except where indicated, Figures 2C and 2D). Flies were transferred to fresh media every three to four days, except for the experiment which controlled for ethanol evaporation by transferring every day (Figure 2C).

### Ethanol concentrations of fly diets

Evaporation and bacterial metabolism may decrease the effective ethanol concentration of the fly diets. Using a clinical grade breathalyzer, we developed a method to measure ethanol vapor within the headspace of a vial and use this a proxy for dietary ethanol concentration [following (Morton et al. 2014)]. Briefly, a 14-gauge blunt needle attached to 50 mL syringe is used to sample the headspace of vial. The sampled air is then pushed through the mouthpiece of an Intoximeters Alco-Sensor® III breathalyzer. Using 2.5%, 5%, and 10% ethanol media, with either 20 bacterially-colonized or bacteria-free flies, we checked ethanol concentration once per day for four days. Four (2.5% and 5%) or five (10%) replicate vials of each of the ethanol treatments were used. Preliminary experiments show that ethanol vapor concentration in the headspace stabilizes within two hours of opening a vial or taking a measurement (data not shown).

### Inebriation Assay

Inebriation was measured using an established method (Sandhu et al. 2015). Briefly, vials were gently tapped and the number of individuals that were able to stand up 30 seconds later was recorded. Inebriation was measured on bacterially-colonized and bacteria-free flies on diets containing 5%, 10%, 12.5% and 15% ethanol, with four independent replicates per ethanol and bacterial treatment. As a positive control, one mL of 85% ethanol was added to a cellulose acetate plug that was pushed into the middle of a vial, and this vial was capped tightly with a rubber stopper. Within 30 minutes, this method leads to inebriation in approximately 50% of flies under a variety of experimental conditions (Sandhu et al. 2015). To measure ethanol vapor in the positive control, a valve was attached to the rubber stopper and the headspace was sampled at 30 minutes using the breathalyzer method described above.

### Internal ethanol concentration of flies

To quantify the ethanol concentration to which fly tissues are exposed, we measured the internal ethanol concentration using a colorimetric enzymatic assay (Sigma-Aldrich MAK076). This approach measures the combined effects of ethanol uptake and internal metabolism. We measured ethanol concentration in individual flies fed 0% or 10% ethanol diets for 15 days. As a positive control, a group of flies not previously exposed to ethanol were enclosed in a rubber-stoppered vial with a cotton ball soaked with two ml of 35% ethanol [similar to (Fry 2014)]. A dry cotton ball was added above the ethanol soaked one so that flies were unable to ingest ethanol, while still being exposed to ethanol vapor. After 60 minutes, individuals that could not stand, but still showed leg movements were selected. To calculate final internal concentration per fly, ethanol was considered to be primarily located in the hemolymph and the hemolymph volume was assumed to be 85 *μ*L per fly (Troutwine et al. 2016; Cowmeadow et al. 2005).

### Measurement of fecundity and lifespan

Lifespan and fecundity were measured simultaneously during the same experiment. Four replicate vials of 20 females each were created for the two bacterial treatments (bacterially-colonized and bacteria-free) and the seven ethanol treatments (0% to 15%, in 2.5% increments) resulting in 56 total vials for the 14 treatments. Survival was checked each day and dead flies were removed with each transfer. Fecundity was calculated by the number of adults that emerge per transfer to new diet, divided by the number of females alive at the start of that transfer, summed over the entire experiment. Approximately 90% of all pupae that formed survived to adulthood with no differences in eclosion rate between ethanol or microbial treatments (Figure S6) and thus only adult emergence data is shown. Development rate was measured as the day the first pupae formed following a transfer to a new vial. In a follow-up fecundity experiment that controlled for ethanol evaporation, flies were transferred to new freshly-inoculated media every day. This experiment also included a diet that was isocaloric with the 2.5% ethanol diet. The isocaloric diet was created by the addition of 4.4% glucose (to the 10% glucose added to all diets) and assumes ethanol is 7 kcal/g and glucose is 4 kcal/g (Ja et al. 2007).

### Bacterial abundance within flies

This experiment was set up identical to the lifespan and fecundity experiment, except that only three replicate vials were used. On days 14, 21, 28, and 31, one to three individual flies from each replication and treatment were externally sterilized, homogenized, serially diluted, and plated onto MRS media (Obadia et al. 2017). For the 12.5% and 15% ethanol treatments, we did not sample flies on days 31, and 28 and 31, respectively, because of fly death before the end of the experiment. Eight to 16 individuals were plated per ethanol treatment (mean=11.5). Colony forming units (CFUs) were identified by visual comparison to laboratory stocks of various species of *Acetobacter* and *Lactobacillus.* Additionally, the identity of representative CFUs was confirmed using 16S rRNA sequencing (Supplementary Data X). In only one of 81 individual flies sampled was there a CFU that had neither *Acetobacter* nor *Lactobacillus* morphology. Because this CFU morphology represented less than 2% of the total bacterial community of this fly, it was disregarded as potential contamination.

### Bacterial sensitivity to ethanol in vitro

We tested *A. pasteurianus*, *L. plantarum*, and *L. brevis*, isolated in the bacterial abundance experiment, for sensitivity to ethanol. Isolates were grown overnight at 30°C in an appropriate medium (MYPL for *A. pasteurianus* and MRS for *L. plantarum* and *L. brevis*) and diluted to a working OD of 0.01. For *A. pasteurianus* and *L. plantarum*, growth was measured in 0% to 15% ethanol media in a 96-well plate using a TECAN Infinite F200 PRO, set to 30°C and 5 minutes of orbital shaking per 10 minutes. For *L. brevis*, which forms a pellet when grown in a 96-well plate, two mL of 0% to 15% ethanol MRS was inoculated with the overnight culture and shaken continuously in cell culture tubes at 30°C. After 24 hours, maximum final OD was determined for each isolate, and a two-parameter Weibull function was fit to the normalized maximum ODs from the aggregate data for each strain (R package drc: Analysis of Dose-Response Curves). The inhibitory concentration for 50% growth (IC50) was calculated as the ethanol percentage that reduced normalized maximum OD by half.

### Bacterial abundance on the diet

Experiments were set up as above, except only 0%, 7.5%, 10%, 12.5%, and 15% ethanol diets were used. On day three five flies from each of four replicates per treatment were individually homogenized, serially diluted in a 96-well plate, and pinned on selective media (MRS for *L. plantarum*, MRS+X-Gal for *L. brevis*, and MYPL for *A. pasteurianus*) using a 96-pin replicator (Boekel), (Obadia et al., 2017 and/or Gould et al., 2018). After fly removal from the vials, one mL of PBS and approximately ten glass beads were added. This was shaken gently on a Nutator at speed 3 for ten minutes, at which time 200 *μ*L was serially diluted and pinned as above. To convert pinned colony growth to actual bacterial abundance, overnight cultures of *L. plantarum, L. brevis*, and *A. pasteurianus* were serially diluted in 96-well plates as above. These serial dilutions were both plated onto agar plates (to determine actual abundance) and pinned (to determine pinning efficacy). A standard curve was created relating actual abundance to growth due to pinning.

### Intestinal Barrier Failure

We measured the level of intestinal barrier failure (IBF) by supplementing fly diet with 2.5% (w/v) FD&C Blue No. 1 (Rera et al. 2012). Two independent experiments were done, the first with 0% and 5% ethanol diets and the second 0%, 5%, and 7.5% ethanol diets, each with bacterially-colonized and bacteria-free treatments. For each, three or four vials of 10 flies were monitored over their entire lifespan and degree of IBF determined by the amount of blue coloration in tissues upon death. For statistical purposes, individuals in IBF categories 0 and 1 were considered IBF negative and individuals with IBF categories 2 and 3 were considered IBF positive (Clark et al. 2015). No significant differences were found between experiment 1 and experiment 2, so they were combined into a single dataset. Because the blue dye accumulates in flies with IBF and increases mortality (Clark et al. 2015), we did not directly compare the lifespan data from these IBF experiments with experiments lacking blue dye.

### Lipid content

Bacterially-colonized or bacteria-free flies were reared on 0%, 5%, and 10% ethanol diets for 16 days, as described above. Four to ten individuals were pooled by sex (mean=9.5), with three to five replicates for each bacteria-ethanol-sex treatment. The mass of pooled flies was determined to the nearest 1/10 of a milligram on a Mettler Toledo microbalance. Free and total lipid content was determined using established colorimetric methods (SIGMA F6428, T2449, and G7793), (Wong et al. 2014; Tennessen et al. 2014).

### Measurement of gene expression

We used NanoStrings profiling to quantify *D. melanogaster* gene expression changes due to ethanol ingestion and bacterial colonization (NanoStrings Technologies, Inc. Seattle, WA, USA). A custom NanoStrings probeset was designed to target genes related to ethanol metabolism, innate immunity and inflammation, ethanol-mediated behavior, among others (A full list of genes, raw counts, normalized counts, and P-Values are found in Supplementary Dataset S1). Additionally, probes were designed to the bacterial 16S ribosomal RNA and alcohol dehydrogenase A and B genes. Bacterially-colonized or bacteria-free flies were reared on 0% and 10% ethanol diets for 11 days. Total RNA was obtained from individual whole flies using a Trizol/Chloroform extraction [following (Elya et al. 2016)]. 50 to 75 ng of purified RNA per sample was hybridized to the NanoString reporter and capture probesets following manufacturing instructions, and profiled on an nCounter SPRINT machine (Laboratory of Greg Barton, UC Berkeley). Raw counts were normalized to internal NanoStrings positive and negative control probes and three housekeeping genes (Actin 5C, Gadph, and Ribosomal Protein L32). The correlation between each of the three housekeeping genes and the final normalization factor was always greater than 0.89. Treatment effects were determined with a two-way ANOVA using ethanol and bacterial colonization as independent variables and normalized counts (i.e. expression level) as the dependent variable (Supplementary Dataset S1). Significance was determined using a 5% Benjamini-Hochberg False Discovery Rate. For all bacterial16S genes, the normalized counts were greater than 10-fold higher in the bacterially-colonized treatments compared to the bacteria-free treatments (Supplementary Dataset S1). For all bacterial ADH genes, the normalized counts in the bacteria-free treatment were within three standard deviations of the negative control probes and were greater in the bacterially-colonized treatments (Supplementary Dataset S1).

## Results and Discussion

### Establishing a model for chronic ethanol ingestion in flies

Here we investigated the effects of chronic ethanol ingestion on *D. melanogaster* adults and if the microbiome can mediate these effects. We first developed an administration method in which ethanol is ingested directly from the media and asked how this method compares to established methods of ethanol administration in adult flies. Previous methods used liquid ethanol added to tightly capped vials, causing flies to absorb ethanol vapor through their cuticle.

For all subsequent experiments, we used two microbiome treatments: Bacteria-free and bacterially-colonized. Bacteria-free flies were generated using established protocols (Koyle et al. 2016) and bacterially-colonized flies were created by allowing approximately 50 normally-colonized adults (from unmanipulated lab stocks) to seed autoclaved media with their frass, removing these flies, and then introducing bacteria-free flies.

We first wanted to know how ethanol headspace vapor in our experiments compares with the vapor levels of ethanol used in established ethanol inebriation studies and if headspace vapor serves as a good proxy for dietary ethanol. To measure ethanol content, we developed a low-cost and rapid method to measure ethanol in the vapor headspace of the fly vial using a breathalyzer and used this as a proxy for dietary ethanol content [following (Morton et al. 2014)]. We sampled the headspace vapor of freshly prepared vials with 0% to 15% dietary ethanol added. Additionally, we used two methods to expose flies to ethanol vapor. In the first, we soaked a cotton ball with 2 mL of 35% ethanol and covered with a dry cotton ball so flies could not ingest the ethanol [similar to (Fry 2014)]. In the second, we added 1 mL of 85% ethanol to a cellulose acetate plug (Sandhu et al 2015). We found that headspace vapor accurately measures dietary ethanol (Figure 1A and S1). We also found that these methods lead to ethanol vapor levels many times greater than our dietary ethanol method (Figure 1A). Therefore, our chronic ingestion model exposes flies to a much lower headspace vapor than previously established acute inebriation models, and suggests the main source of ethanol uptake is ingestion.

To confirm that flies effectively uptake ethanol when it is mixed directly in the media, we measured the internal ethanol concentrations of flies fed ethanol diets. We found that flies fed 10% ethanol diets contain higher internal concentrations of ethanol than flies fed 0% ethanol diets, which shows that in our treatment conditions flies successfully ingest dietary ethanol (Figure 1B). We were also interested in how internal ethanol concentrations in flies fed ethanol compare to flies exposed to ethanol vapor. We found that inebriated flies (exposed to 35% ethanol vapor, which causes most flies to become immobile within an hour) have even higher internal ethanol levels. This suggests that our dietary regime exposes flies to sub-inebriating levels of ethanol.

We next asked if flies show behavioral signs of intoxication using the inebriation assay of Sandhu et al. 2015, which measures inebriation as the inability for flies to stand after gently tapping the vials. We found that, after 30 minutes, less than 5% of flies show signs of inebriation, even on the highest ethanol diets, while half of flies exposed to 85% ethanol vapor are inebriated (Figure 1C). Importantly, there was no effect of bacterial treatment on inebriation, consistent with results from a previous study that used antibiotics to clear flies of their bacterial communities (Sandhu et al 2015).

Finally, we asked how dietary ethanol concentration changes over time and hypothesized that both evaporation and bacterial metabolism reduce ethanol content. As expected, the ethanol vapor decreases over time (Figures 1D-1E). For 10% ethanol media, approximately half the ethanol remained after 3 days. Colonization of vials with fly gut bacteria reduced the ethanol levels further, particularly at low concentrations. For example, in the 2.5% ethanol treatment, there is no detectable ethanol in the bacterially-colonized treatment on day two, but for the 10% ethanol treatment there was no difference in the ethanol concentrations until day four. These results suggest ethanol loss by two mechanisms. First, evaporation decreases ethanol concentration. Second, bacterial metabolism consumes ethanol. The lag in bacterial ethanol degradation presumably occurs because bacterial populations in the vials start off small. All vials are initially sterile and are inoculated with the transfer of bacterially-colonized flies. Thus bacterial abundance only becomes great enough to affect the measured ethanol on day two or later.

Taken together, we have established an experimental model of chronic ethanol ingestion in adult *D. melanogaster*. Ethanol remains in the media long enough for flies to uptake, ethanol is detectable internally after ingestion, and flies do not show overt signs of inebriation. By adding ethanol directly to the diet, we mimicked the route of natural ingestion for flies and increase the translational power of our model, as humans consume ethanol via their diet rather than through inhalation.

**Figure 1:**
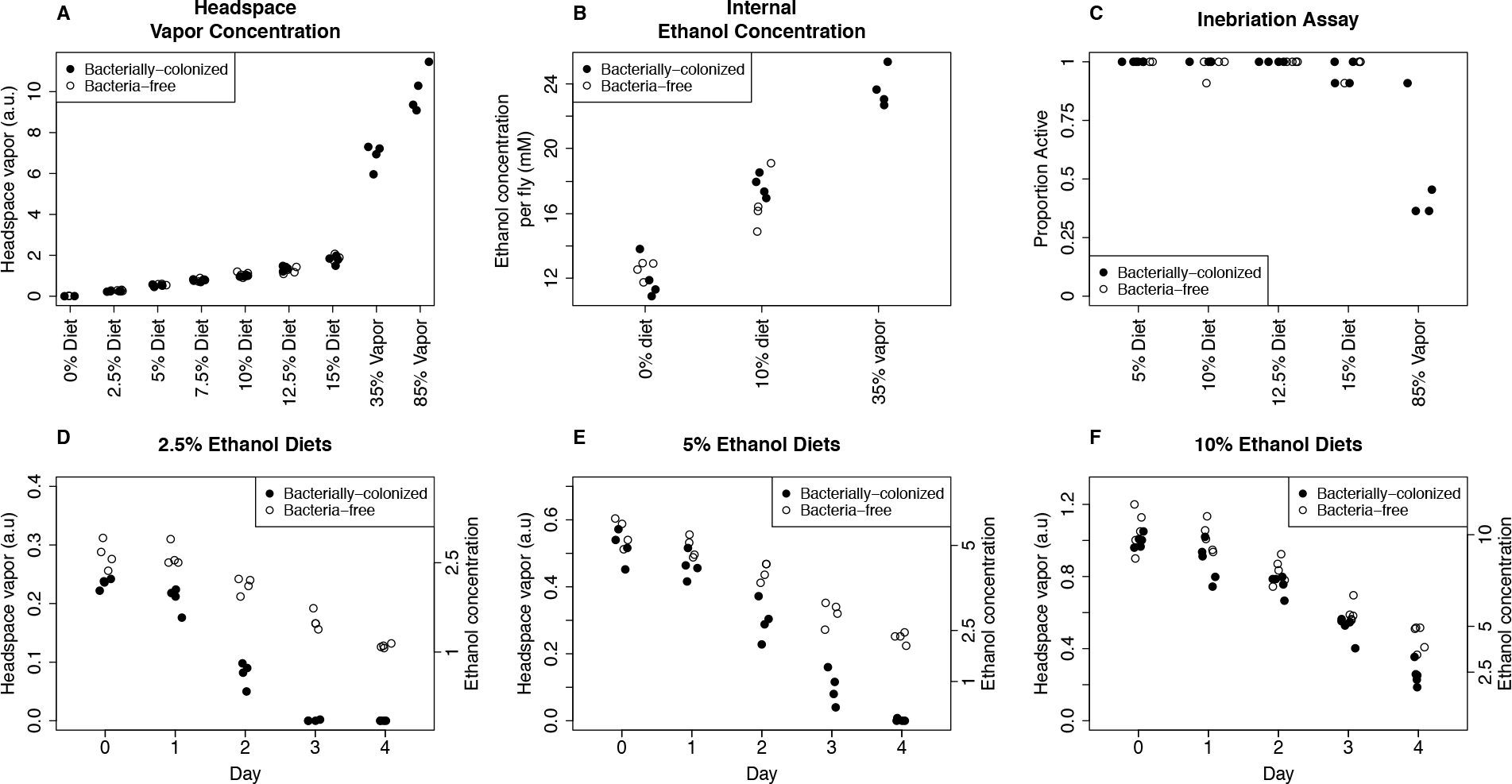
Establishing a chronic ethanol ingestion model in *Drosophila*. **A: Dietary ethanol leads to less headspace ethanol vapor than standard methods for ethanol inebriation.** Headspace vapor was determined by sampling the vial headspace with a syringe and forcing this mixture through a medical-grade breathalyzer. For the 35% and 85% ethanol treatments, either a cotton ball (35%) or a cellulose acetate plug (85%) was soaked with liquid ethanol in a tightly capped vial (Fry 2014; Sandhu et al. 2015). **B: Flies uptake ethanol from their diet, but this still leads to lower internal ethanol concentrations than inebriating levels of ethanol vapor.** Internal ethanol concentration was assayed enzymatically on individual flies fed either 0% or 10% ethanol diets or exposed to 35% ethanol vapor (Fry 2014). **C: Dietary ethanol does not lead to inebriation by standard assay.** Proportion active is the proportion of 11 individual flies that can stand up after gently tapping the vial 30 minutes after initial exposure to ethanol. 85% ethanol vapor (final column) robustly leads to inebriation in about half of the individuals at this timepoint (Sandhu et al. 2015). **D, E and F: Dietary ethanol content decreases over time with greater loss in bacterially-colonized treatments.** Ethanol concentration (right axis) was calculated from day 0 measurements (Figure S1). Measurements from 0% ethanol media are always below 0.02 and are therefore not shown. Note that flies are transferred to fresh vials on day 3 or 4 (Figure 2A and B, 3, 4, and 5) or day 1 (Figure 2C).

### Bacterial colonization of flies masks the negative effects of ethanol on lifespan

We sought to determine the effects of ethanol and the microbiome on fly fitness, focusing on lifespan and fecundity (Gould et al. 2018), which have not been investigated in a fly ethanol model. While the results in humans are conflicting at very low ethanol consumption, in general ethanol consumption is associated with shorter lifespan (Wood et al. 2018). The natural habitat of *D. melanogaster*, fermenting fruit, often contains 1-5% ethanol and the unnatural but common habitat of vineyards can contain up to 10% ethanol (Gibson et al. 1981). We therefore tested dietary ethanol concentrations from 0% to 15%, which spans from ecologically relevant concentrations to concentrations above those to which flies are normally exposed.

We measured lifespan, fecundity, and microbiome composition (see next section) in the same experiment. Four replicate vials of 20 flies were used for each ethanol and bacterial treatment. In our first experiment, we transferred flies to fresh food every three to four days to balance between maintaining dietary ethanol concentration, which decreases over time, (see Figure 1D-F) and maintaining bacterial colonization, which requires less frequent transfers (Blum et al. 2013)].

Bacterially-colonized flies consistently showed a shorter lifespan than bacteria-free flies, in agreement with previous studies (Figure 2A; Table S1; Data for individual flies is shown in Figure S2; Lifespan curves are shown in Figure S3), (Ridley et al. 2012; Clark et al. 2015; Steinfeld 1927). However, the shorter lifespan in bacterially-colonized flies was robust to dietary ethanol, with no significant ethanol-induced decrease in average or maximum lifespan observed except at levels above those experienced in nature (12.5% and 15%), This was in sharp contrast to the bacteria-free flies, which overall live longer, but show a nearly linear and dose-dependent decrease in average and maximum lifespan beginning at just 2.5% ethanol. Overall, these data suggest that two independent mechanisms interact to determine lifespan in this system: bacterial colonization and ethanol exposure. First, the effect of bacterial colonization is dominant to the effect of ethanol at levels below 10%. Second, there is a clear negative effect of ethanol, but its effect is completely masked by bacteria at low to moderate ethanol concentrations.

That bacterially-colonized flies have a reduced lifespan compared to bacteria-free flies has been reported before though never with the same magnitude we found here, suggesting that the flies in our lab may be colonized with a particularly lifespan-shortening consortium of bacteria. However, an equally plausible explanation would be that our media, which lacks the commonly-used microbial growth inhibitor tegosept, may have greater bacterial loads than other studies (Obadia et al. 2018). This would lead to a greater difference in dietary bacterial load between the bacterially-colonized and bacteria-free treatments which could explain the more drastic lifespan reduction that we observe.

**Figure 2:**
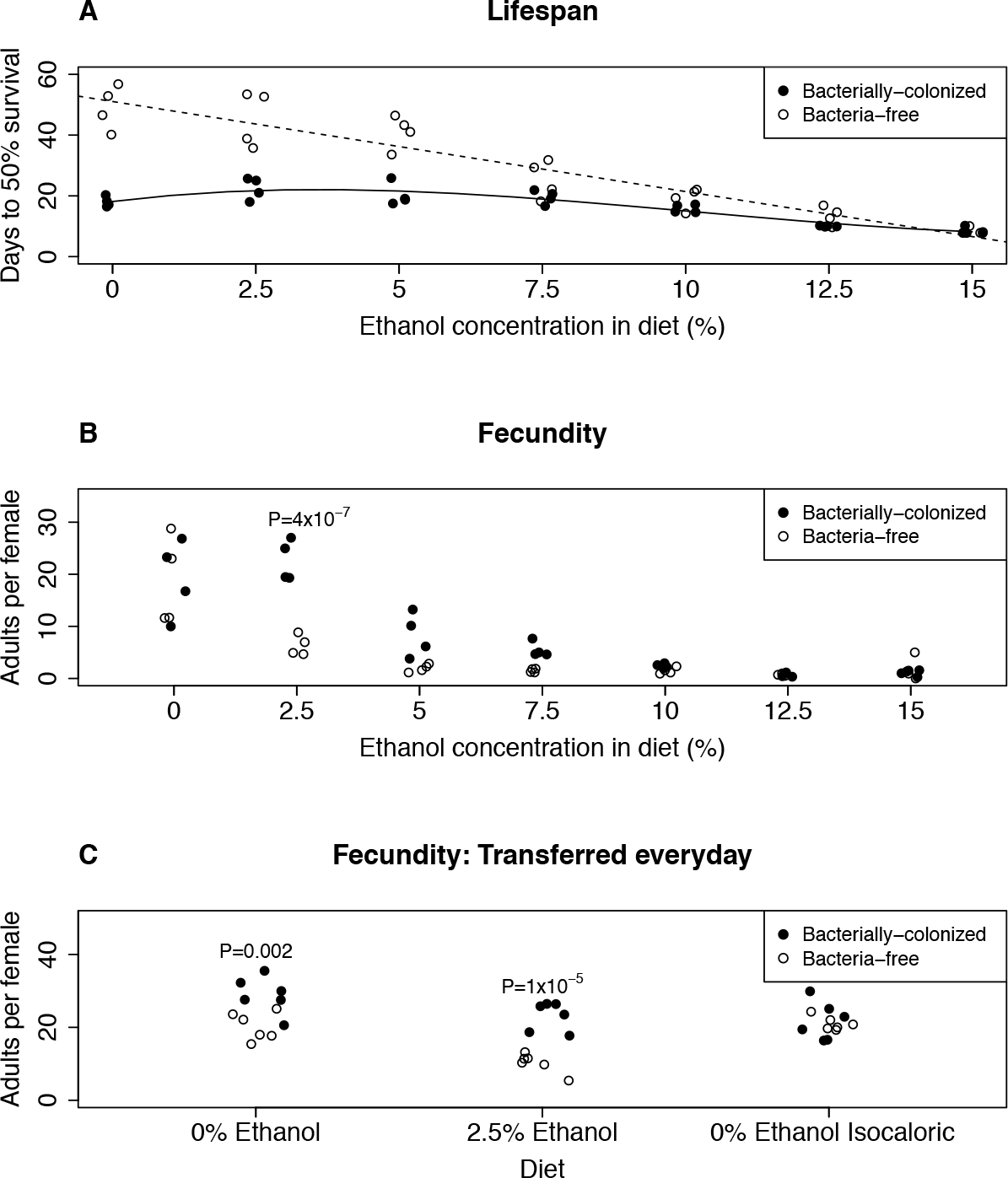
Bacteria mediate the effect of ethanol on fly fitness. **A: Bacterial colonization and ethanol negatively affect fly lifespan, with the negative effect of ethanol being unmasked in bacteria-free flies.** Days to 50% survival is per replicate and calculated from birth (see methods). Each replicate began with 20 female flies. Lines show the best fit lines for each bacterial treatment (Solid: bacteria-free, linear, R^2^=0.874; Dashed: bacterially-colonized, third order, R^2^=0.832). Data for individual flies is shown in Figure S2 and Table S1. **B: Bacteria ameliorate the negative effects of ethanol on fly fecundity.** Adults per female is calculated by the number of adults that emerge per flip, divided by the number of females alive at the start of the egg laying period. Each replicate began with 20 females. P-values are calculated from a pairwise t test between bacterial treatments for a given ethanol treatment and are Holm-Bonferroni corrected for multiple comparisons. Non-significant P-values are not shown. An independent replication of this experiment (with 0% and 2.5% ethanol) is shown in Figure S4. **C: The effect of 2.5% ethanol on fecundity is not due to ethanol evaporation or the caloric contribution of ethanol.** Flies were transferred to fresh diets each day to reduce the effect of ethanol evaporation (Figure 1D). The isocaloric diets have added glucose so they contain identical calories as the 2.5% ethanol diets. Adults per female was calculated as in Figure 2B, however in this experiment each replicate began with 10 females per replicate. P-values are calculated from a pairwise t test between bacterial treatments for a given dietary treatment and are Holm-Bonferroni corrected for multiple comparisons. Non-significant P-values are not shown.

### Bacteria ameliorate the negative effects of ethanol on fecundity

We found a strong effect of ethanol on fly fecundity that is mediated by bacterial treatment (Figure 2B). Without ethanol, there was no difference in fecundity between bacterially-colonized and bacteria-free flies, which is consistent with previous work (Ridley et al. 2012). For both bacterial treatments, ethanol reduced fecundity, but bacteria-free flies were more sensitive: at 2.5% ethanol, bacterially-colonized flies had significantly higher fecundity (P=4×10^−7^, Figure 2B; P=0.03, Figure S4). The same trend was observed on both 5% and 7.5% ethanol diets (though not statistically significant at P<0.05 after correction for multiple comparisons). Interestingly, we found that ethanol does not lead to the typical tradeoff between lifespan and fecundity observed by varying nutrients (Zera & Harshman 2001; Djawdan et al. 1996) ‒ instead we found that ethanol decreases both components of fitness (Figures 2A and 2B, Figure S5). This suggests that ethanol, even at the low concentrations used in this study, is acting more like a toxin than a source of calories.

On 2.5% ethanol media, ethanol content is significantly reduced by day two in the bacterially-colonized treatment (Figure 1D). Therefore, the difference in fecundity observed in the 2.5% ethanol treatment (Figure 2B) could simply be due to less dietary ethanol in the bacterially-colonized treatment. Although the fecundity difference between bacterially-colonized and bacteria-free flies remains even when accounting for differential ethanol loss (Supplementary Dataset 2), we nonetheless repeated the fecundity experiment but transferred the flies to fresh diets every day. To ensure the persistence of the intestinal bacterial communities, we seeded each daily batch of media with the frass of bacterially-colonized flies. Also, to test whether the calories added by the ethanol in the 2.5% treatment affect the flies, we added a 0% ethanol treatment that is isocaloric with the 2.5% ethanol treatment.

The daily transfer experiments showed comparable results. In concordance with the 3-4 day transfers, bacterially-colonized flies had greater fecundity in the 2.5% ethanol treatment relative to bacteria-free flies (P=1×10^−5^, Figure 2C). This strongly suggests that bacterial metabolism of ethanol on the food does not cause the difference in fecundity between the bacterial treatments. We also found no effect of bacterial treatment in the isocaloric diets suggesting that the differences in fecundity cannot be attributed to ethanol’s caloric contribution.

The observed fecundity differences could be due either to maternal egg production or larval survival. To differentiate between these causes, we measured larval development time as a proxy for larval survival because we could not directly count egg laying (and thus could not calculate survival from egg to adulthood). Although development time increased on the highest ethanol diets (Figure S6), we found no effect on development time between bacterially-colonized and bacteria-free treatments at the 0%, and 2.5% ethanol treatments (all pairwise t-tests, P>0.2), consistent with a previous result that ethanol does not affect larval development except in the final larval stage at 5 days (McClure et al. 2011), when most of the ethanol has evaporated from the media (Figures 1D,1E and 1F). Furthermore, previous work has shown that 12% ethanol over the entire developmental period reduces larval survival to 25% (McClure et al. 2011). We find that fecundity drops to near zero at 5% ethanol for bacteria-free flies versus 10% ethanol for bacterially-colonized flies. Thus, maternal egg production, rather than larval survival, accounts for fecundity effects seen in Figure 2B.

Taken together, these results suggest significant ecological and evolutionary impacts of microbes in mediating the negative effects of ethanol toxicity on fly fecundity. While the exact doses of ethanol that flies consume in the wild remains obscure, the concentration in naturally fermenting fruit is typically 1-5% and can be as high as 10% in wineries (Gibson et al. 1981). In all cases for flies fed 2.5% to 7.5% ethanol, we found that bacterially-colonized flies had higher fecundity than bacteria-free flies and this effect persists even when controlling for ethanol metabolism by daily transfers to fresh media. Thus, at ecologically relevant concentrations of ethanol, bacterial colonization mitigates the negative effects on fecundity.

### Ethanol shifts the composition of bacteria associated with D. melanogaster

Diet is a strong determinant of microbiome composition in flies and other animals. In particular, fruit feeding flies, which are exposed to naturally produced dietary ethanol, have significantly different bacterial and yeast communities than flies collected from other substrates (Chandler et al. 2011; Chandler et al. 2012). We hypothesized that the bacterial communities associated with flies would shift in response to ethanol ingestion. In particular, we expected that ethanol would strongly decrease the total abundance of bacteria in high ethanol treatments and these shifts would favor the abundance of bacteria with low sensitivity to ethanol. Thus, in a parallel replicate of the lifespan-fecundity experiment (Figures 2A and 2B), we determined fly bacterial load and composition by homogenizing individual flies and plating onto selective media. The different bacterial strains were identified by colony morphology.

We found that total bacterial load per fly was between 9×10^3^ and 3×10^6^ colony forming units (CFUs) for the 0% ethanol containing diets (mean=7×10^5^). This is comparable to previous studies of *D. melanogaster* (Blum et al. 2013; Obadia et al. 2017). Contrary to our expectations, we found that total bacterial load was relatively constant up to the highest ethanol treatment (Figure 3A). We next asked how the bacterial composition changes in response to ethanol. In agreement with the previous work in our laboratory and that of others, our flies are dominated by species in the genera *Acetobacter* and *Lactobacillus* (Broderick & Lemaitre 2012). Different bacteria had different responses to dietary ethanol. *Acetobacter pasteurianus* concentrations decreased 10-fold from 0% to 2.5% ethanol and remained constant until 12.5% ethanol where they dropped to essentially 0 (Figure 3B). Conversely, we found that the response of the *Lactobacilli* to ethanol was remarkably different than *A. pasteurianus*. The abundance of *L. brevis* increased with dietary ethanol and this was the only species that was present in all flies at 15% ethanol (Figure 3C). *L. plantarum* was most abundant at intermediate concentrations of ethanol, but like *L. brevis*, it did not appear as sensitive to high levels of ethanol as *A. pasteurianus* (Figure 3D).

To confirm the direct effect of ethanol on the bacterial growth, we measured the *in vitro* growth response to ethanol of *A. pasteurianus*, *L. plantarum* and *L. brevis* strains isolated during the experiment in Figure 3A-3D. These experiments confirmed that *A. pasteurianus* is more sensitive to ethanol than *L. brevis* and *L. plantarum* (Figure 3E). These results indicate that the bacterial composition of flies varies, at least in part, according the ethanol sensitivities of the bacterial strains. However, because these *in vitro* experiments show that these bacteria are more sensitive to ethanol than is suggested by their *in vivo* abundances, we hypothesized that the fly intestine protects bacteria from ethanol toxicity. In support of this, we found high abundance of *L. brevis* and *L. plantarum* within flies fed a 15% ethanol diet despite these bacteria being undetectable on this media (Figure S8). Similarly, *A. pasteurianus* is present within flies fed 12.5% ethanol despite this bacterium being absent on this media (Figure S8). This confirms that the host shields the effect of ethanol on the bacteria.

**Figure 3:**
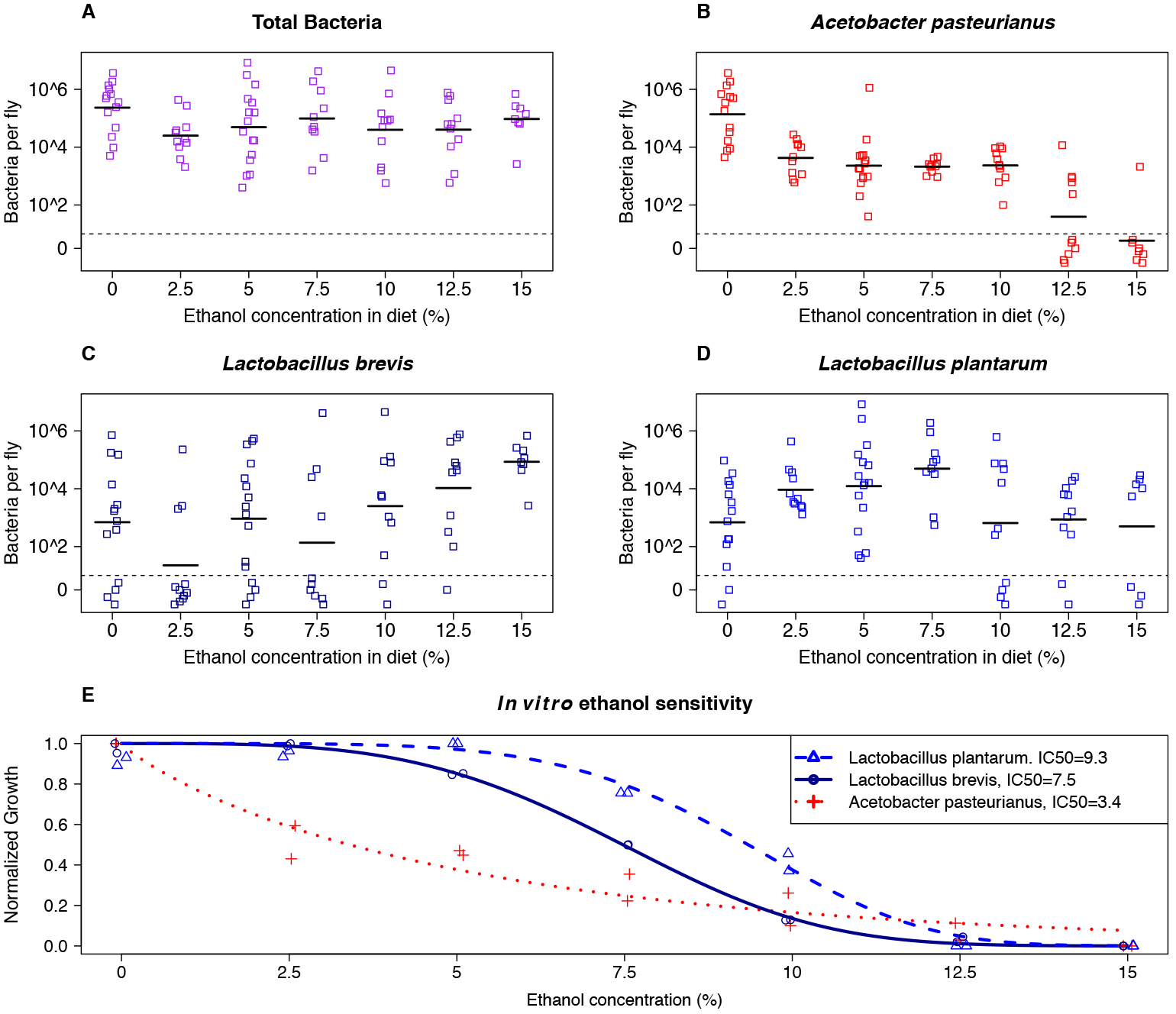
Bacterial community dynamics in response to ethanol diets. **A, B, C and D: The abundance of *Acetobacter pasteurianus* decreases with increasing dietary ethanol, while the abundance of *Lactobacillus plantarum* and *Lactobacillus brevis* remain high.** Each point represents an individual fly. All points below the dashed line are 0 and are expanded for clarity. The black bars represent the mean of the log transformed bacterial load. Number of individual flies per treatment: 0% N=14; 2.5% N=11; 5% N=16; 7.5%N=10; 10%N=11; 12.5% N=11; 15%N=8. We found no effect of fly age [multivariate ANOVA (Adonis, package vegan in R; P = 0.159)] and therefore all four timepoints are pooled (see methods). **E. *A. pasteurianus* is more sensitive to ethanol than *L. plantarum* or *L. brevis in vitro*.** Strains were isolated in the *in vivo* bacterial abundance experiment (Figure 4). Growth was measured using MRS or MYPL liquid media containing 0% to 15% ethanol in either a 96-well plate (*A. pasteurianus* and *L. plantarum*) or cell culture tubes (*L. brevis*) for 24 hours at 30C. Datapoints are the final normalized OD of two independent replicates. A two-parameter Weibull function was fit to the normalized ODs from the aggregate data for each strain (R package drc: Analysis of Dose-Response Curves). The inhibitory concentration for 50% growth (IC50) was calculated as the ethanol percentage that reduced normalized maximum OD by half.

### Dietary ethanol decreases intestinal barrier failure

To explore the fly physiology underlying mortality following ethanol ingestion in flies, we examined intestinal barrier failure (IBF), which is strongly linked to alcoholic liver disease in humans (Chen & Schnabl 2014) and is a hallmark of aging-related death in flies (Rera et al. 2012; Clark et al. 2015). We used the Smurf assay (Rera et al. 2012), rearing flies on a diet containing blue dye no. 1 and scoring them for a blue body coloration, which is indicative of a permeabilized gut. Consistent with previous results (Rera et al. 2012; Clark et al. 2015), we found that nearly all flies on a 0% ethanol diet show IBF upon death. Quite surprisingly, on ethanol diets, we found a significant decrease in the proportion of flies that show IBF (Figures 4A and S9). Furthermore, bacteria-free flies (which are more sensitive to ethanol) show significantly less IBF than bacterially-colonized flies on ethanol diets. Examining the effects of bacteria and ethanol together, we detect a significant interaction (P=1×10^−4^, Figure 4A). Taken together, this suggests that IBF is not the causative mechanism of ethanol-induced lifespan decline in flies. That some individuals in the ethanol treatments nonetheless showed IBF can be explained by the normal background aging process (Rera et al. 2012; Clark et al. 2015).

**Figure 4:**
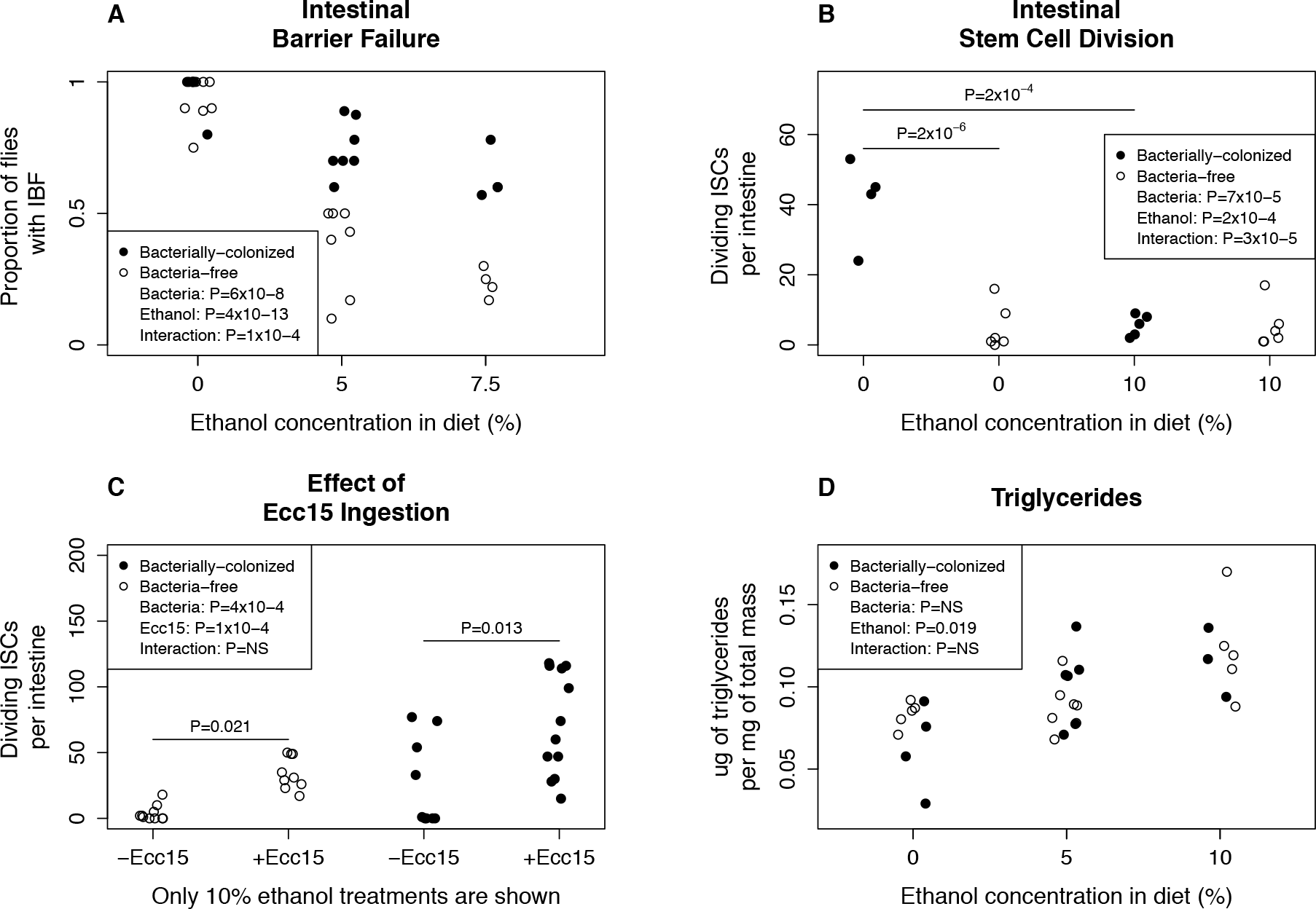
Bacteria mediate the effect of ethanol on fly physiology. Within each panel, values in the legend are the results from a two-way ANOVA. Comparison P values are calculated from a pairwise t test between treatments and are Holm-Bonferroni corrected for multiple comparisons within an experiment. All data shown is for females. Male data for IBF is shown in Figure S9 and for triglyceride content is shown in Figure S11. **A. The prevalence of intestinal barrier failure (IBF) decreases with dietary ethanol and this decrease is greater in bacteria free flies.** Each point represents the average from a replicate vial. Flies were scored within 24 hours of death. **B. Ethanol reduces intestinal stem cell turnover in bacterially colonized, but not bacteria free, flies.** Each datapoint indicates the number of pH3 stained cells in an individual intestine. Results from an independent experiment as shown in Figure S10. **C: Ethanol does not inhibit the ability of ISCs to regenerate following oral ingestion of *Erwinia carotovora carotovora 15* (Ecc15).** Each datapoint indicates the number of pH3 stained cells in an individual intestine. **D: Ethanol ingestion increases stored triglycerides in flies, regardless of bacterial treatment.** Each point represents a pooled sample of 4 to 10 flies.

### The effect of ethanol on ISC turnover is microbiome dependent

Intestinal barrier function is maintained through controlled intestinal stem cell (ISC) turnover (Lemaitre & Miguel-Aliaga 2013). Because stem cell hyper-proliferation leads to loss of intestinal function (Li & Jasper 2016), we hypothesized that the reduction in IBF was due to a decrease in ISC turnover. To quantify stem cell turnover, we measured mitotic cells in the gut by phospho-histone H3 antibody staining (Apidianakis & Rahme 2011). In two independent experiments, we found that in the absence of ethanol, ISC division was significantly greater in bacterially-colonized flies (P=2×10^−6^, Figure 4B; P=2×10^−6^, Figure S10), consistent with previous work showing that ISC hyper-proliferation caused by commensal bacteria shortens fly lifespan [(Figure 2A), (Buchon et al. 2009; Guo et al. 2014)].

In the presence of ethanol, we found that ISC division is significantly decreased in bacterially-colonized flies, but unchanged in bacteria-free flies (P=2×10^−4^, Figure 4B; P=3×10^−6^, Figure S10). These results are in accord with the IBF data presented in Figure 4A for bacterially-colonized flies (specifically, that both IBF and ISC division, two processes linked to intestinal homeostasis, are reduced), but suggest an additional mechanism reduces IBF in bacteria-free flies fed ethanol. Why these two phenotypes are uncoupled in bacteria-free flies, in which ethanol decreases IBF with no change in ISC turnover, remains unknown and suggests different mechanisms of ethanol-induced pathology in bacterially-colonized and bacteria-free flies.

The decrease in dividing ISCs in bacterially-colonized flies led us to hypothesize that ethanol might inhibit the ability of ISCs to regenerate following a biological or chemical challenge. To test this hypothesis, we infected flies with *Erwinia carotovora carotovora 15* (Ecc15), a non-lethal pathogen of *Drosophila*, which reliably induces ISC division following oral ingestion (Buchon, Broderick, Poidevin, et al. 2009). In both bacteria-free and bacterially-colonized flies ingesting ethanol, infection with Ecc15 increases ISC division (P=0.021 and P=0.013, respectively, Figure 4C). Thus, ethanol does not inhibit the ability of ISCs to regenerate despite the observed decrease in ISC division in bacterially-colonized and ethanol-fed flies (Figures 4B and S10).

### Ethanol ingestion increases stored triglycerides in flies

The maintenance of intestinal homeostasis with ethanol treatment may be due to a change in overall fly metabolism. In flies, poor quality diets are linked to both ISC turnover and obesity (Skorupa et al. 2008; Regan et al. 2016). In humans, increased fat deposits in the liver are a hallmark of alcoholic liver disease (Diehl 2002). We hypothesized that ethanol ingestion is leading to greater accumulation of stored triglycerides in flies. Triglycerides are a primary molecule for fat storage in flies and are mainly found in adipocytes within the fat body, an organ analogous to the mammalian liver that is responsible for the majority of energy reserves in adult fly (Arrese & Soulages 2010). We therefore measured stored triglycerides in bacteria-free and bacterially-colonized flies on 0%, 5% or 10% ethanol diets. Consistent with our hypothesis, we found that dietary ethanol increases triglycerides regardless of bacterial colonization, with no effect on either total fly mass or free glycerides (Figures 4D and S11, Table S2). Because dietary sugars increase triglyceride content in flies (Skorupa et al. 2008), our finding is consistent with ethanol acting as an energy source with regards to fat storage, despite the lack of tradeoff between lifespan and fecundity due to ethanol ingestion (Figure S5). The finding that there is no difference in triglyceride content between bacterially-colonized and bacteria-free flies is consistent with the minimal role of bacterial metabolism on 5% and 10% ethanol diets (Figures 1E and 1F) and suggests that fat accumulation does not directly explain either the lifespan (Figure 2A) or intestinal homeostasis (Figures 4A and 4B) results.

### Ethanol ingestion and bacterial colonization affect expression of innate immunity genes

To understand the molecular mechanisms underpinning the differences in lifespan between bacteria-free and bacteria-colonized flies ingesting ethanol (Figure 2A), we surveyed fly gene expression using a custom NanoStrings probeset and selected candidate genes likely to be influenced by ethanol or microbiome status. In concordance with previous work (Broderick et al. 2014), we found many immune system (e.g. lysozyme X and the PGRPs), stress related (e.g. GstD5 and HSP23), and cell differentiation (e.g. upd3) genes to be upregulated in response to bacterial colonization (Table 1). Likewise, and in agreement with Elya et al 2016, we found that anti-microbial peptides (AMPs) as a group show increased expression in bacterially-colonized treatments (Table S3).

We examined many genes and molecular pathways known to mediate the effects of ethanol intoxication in flies, but we found that ethanol only subtle changes the expression of neuropeptideF (Table 1). This is consistent with ethanol ingestion not leading to inebriation (Figure 1C) and flies fed ethanol having a lower internal concentration of ethanol than inebriated flies (Figure 1B).

There are two potential mechanisms that may contribute to the ethanol-induced lifespan reduction in bacteria-free, but not bacterially-colonized, flies. First, host metabolism of ethanol may be more efficient in bacterially-colonized flies, leading to a faster clearance of ingested ethanol. Contrary to this hypothesis, we did not find that genes in the ethanol metabolism pathway [alcohol dehydrogenase (Adh) and acetaldehyde dehydrogenase (Aldh)] were more strongly induced in bacterially-colonized flies (Table 1), which is consistent with the equivalent internal ethanol concentrations we observed for bacteria-free and bacterially-colonized flies fed 10% ethanol (Figure 1B). This result suggests that ethanol metabolism does not directly underpin the differences in lifespan between bacteria-free and bacteria-colonized flies.

In the second mechanism, ethanol may be eliciting the innate immune response in bacteria-free, but not bacterially-colonized, flies. Immune activation promotes shorter lifespan in flies (Garschall & Flatt 2018; DeVeale et al. 2004; Eleftherianos & Castillo 2012) and resistance to ethanol vapor in flies is linked to the innate immunity response (Troutwine et al. 2016). In mammals, dysregulation of the immune response and persistent inflammation is linked to aging (Gomez et al. 2008; Shaw et al. 2010). Two innate immunity genes showed a significant change in expression due to the combination of ethanol and bacterial colonization (Figure 5): Immune deficient (Imd) and Peptidoglycan recognition protein SC1A/B (PGRP-SC1A/B). Imd is a master regulator of innate immunity in response to gram-negative bacteria (Lemaitre & Hoffmann 2007). PGRP-SC1A/B is a peptidoglycan scavenger that negatively regulates the IMD pathway (Kurata 2014). Consistent with their known interaction pathway, PGRP-SC1A/B increased and Imd decreased with ethanol ingestion in bacterially-colonized flies. However, both of these genes show greater changes with ethanol ingestion in the bacterially-colonized treatment, with little or no change in the bacteria-free treatment (Figure 5), and therefore do not directly explain the lifespan differences identified in Figure 2A. We next examined whether antimicrobial peptide (AMP) gene expression, which is a downstream target of Imd, was affected. We found no ethanol dependence of AMP stimulation, (Table S3). This finding is consistent with literature showing that the AMP expression can be muted when Imd is triggered in the absence of a pathogen (Lhocine et al. 2008). Regardless, because of the recognized role of the IMD pathway and innate immunity in regulating fly lifespan, Imd and PGRP-SC1A/B are promising targets to pursue to further our understanding of microbiome by ethanol fitness effects in *Drosophila*.

**Table 1:** Genes showing significant expression changes in response to either ethanol ingestion or bacterial colonization. P-values are adjusted using a 5% Benjamini-Hochberg false discovery rate correction. Genes with less than a 1.5 fold-change or with average normalized counts within two standard deviations of the negative control probes are excluded.

| Genes affected by ethanol ingestion |  |  |
| --- | --- | --- |
| Gene | P-value | Fold-Change |
| neuropeptideF | 0.015 | 1.5 |
| Acetaldehyde Dehydrogenase | 0.035 | 1.6 |
| Acetyl-CoA synthetase | 0.038 | 2.8 |
| HSP70Bc | 0.047 | 3.1 |
| Alcohol Dehydrogenase | 0.047 | 1.8 |

**Table 1: Genes showing significant expression changes in response to either ethanol ingestion or bacterial colonization.**
| Genes affected by bacterial colonization |  |  |
| --- | --- | --- |
| Gene | P-value | Fold-Change |
| PGRP SC1A/B | 0.0029 | 3.3 |
| upd3 | 0.030 | 5.5 |
| LysozymeX | 0.030 | 18 |
| upd2 | 0.030 | 69 |
| Crys | 0.030 | 9 |
| Defensin | 0.030 | 5.3 |
| Charon | 0.030 | 3.8 |
| GstD5 | 0.030 | 2.7 |
| PGRP SD | 0.030 | 4.2 |
| PGRP LB | 0.030 | 2.2 |
| Drosomycin-like 1 | 0.031 | 44 |
| HSP23 | 0.040 | 2.3 |
| PGRP LC | 0.048 | 2.7 |

**Figure 5.**
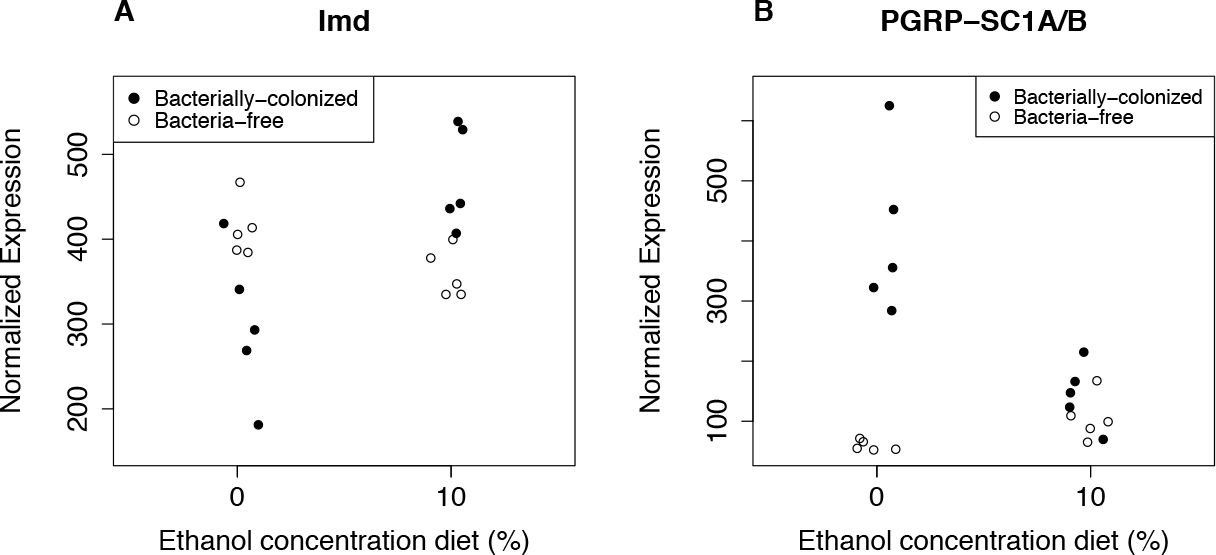
Ethanol induces the Imd response in bacterially-colonized, but not bacteria-free, flies. A two-way ANOVA finds a significant bacteria-by-ethanol interaction on the expression of Imd (Panel A) and PGRP-SC1A/B (Panel B). For both genes, the P-value of the interaction is 0.023 after a 5% Benjamini-Hochberg false discovery rate correction.

### Conclusion: the interaction of bacteria and ethanol shape fly fitness and physiology

Previous studies of the microbiome’s role in alcoholic pathology have focused on alcoholic liver disease, finding that specific bacteria can reduce alcoholic liver disease through a decrease in gut permeability (Forsyth et al. 2009; Bull-Otterson et al. 2013). However, isolating experimental variables has proven challenging due to the complex composition of the microbiome as well as contextual effects that depend on an interaction between the microbiome and diet (Wong et al. 2014).

Our *Drosophila* model of chronic alcoholic pathology shows that ethanol’s effects are mediated by the microbiome. For ecologically relevant levels of dietary ethanol, fecundity was greater in bacterially-colonized flies, highlighting the context-dependence of the microbiome in host physiology. Despite the greater fecundity, bacterially colonized flies had much shorter lifespans than bacteria-free flies, indicating that the microbiome mediates tradeoffs between physiological states (Figure S5).

We were curious to understand how dietary ethanol and the microbiome interact to shape fly lifespan. We propose that in the absence of ethanol or at low ethanol concentrations, the negative effects of bacteria are dominant and reduce lifespan by disrupting intestinal homeostasis and inducing innate immunity (Figure 6). In the absence of a microbiome, the negative effects of ethanol are unmasked, which accounts for the dose-dependent decrease in lifespan of bacteria-free flies. This effect is likely independent of overt inebriation (Figure 1C), disruption of intestinal homeostasis (Figures 4A and 4B), or induction of the innate immune response (Figure 5 and Table S3). We speculate there is an unidentified mechanism of ethanol-induced health decline in flies and this is chiefly observable in bacteria-free conditions.

For bacterially-colonized flies fed higher ethanol concentrations (which have the same lifespan as flies not fed ethanol), two offsetting mechanisms are occurring: First, ethanol is reducing lifespan through the unidentified mechanism described above. Second, ethanol is changing microbiome composition to a less pathogenic state [potentially through the reduction in *Acetobacter* abundance (Figure 3B). Future research will use gnotobiotic flies with defined microbial communities to isolate the independent effects of the microbiome and dietary ethanol.

**Figure 6:**
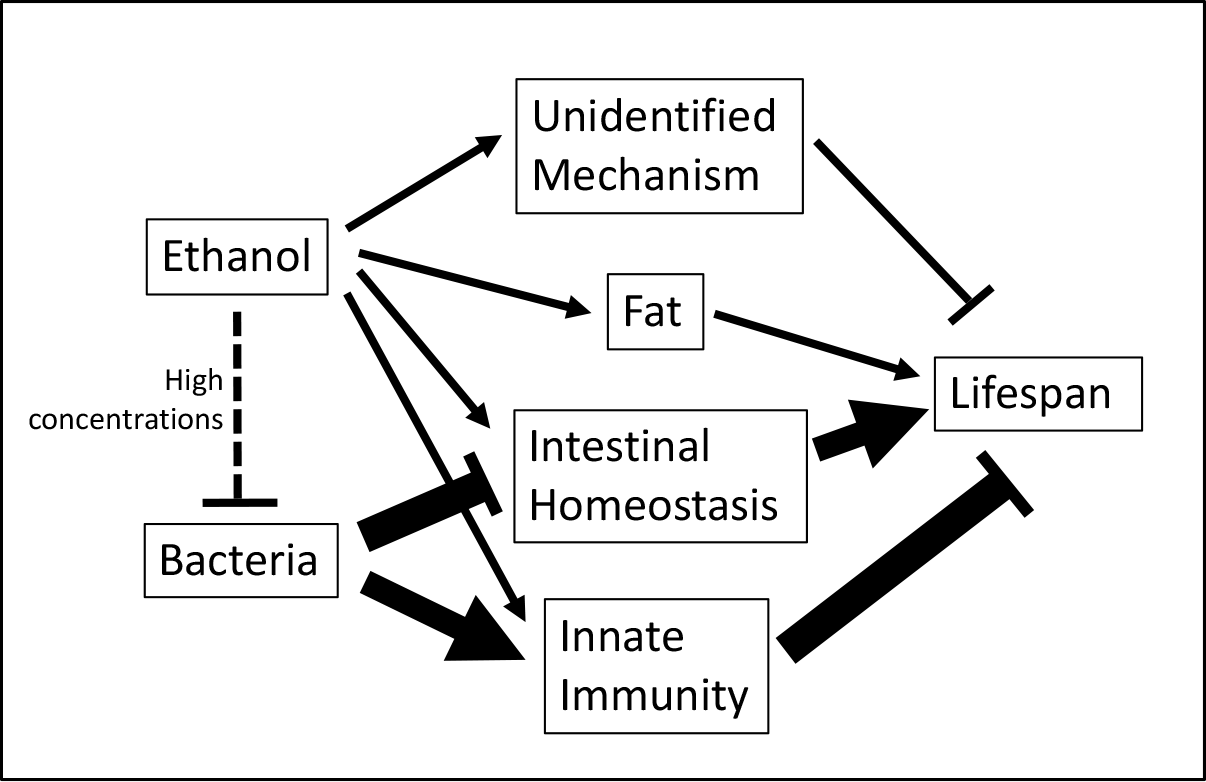
Model of microbiome-dependent ethanol-induced lifespan decline in flies. In the absence of ethanol or at low ethanol concentrations, the negative effects of bacteria are dominant and reduce lifespan by disrupting intestinal homeostasis and inducing innate immunity. At high ethanol concentrations, the microbiome changes composition (primarily through the reduction of *Acetobacter* abundance). This eliminates the bacteria-dependent lifespan reduction, but is offset via an unknown ethanol-dependent mechanism which is damaging to fly health (so the net result is no change in lifespan). This mechanism is primarily independent of intestinal homeostasis and the innate immune response. It is also likely different from known intoxication pathways in flies because our method of ethanol administration does not lead to overt inebriation.

## Acknowledgements

This work is supported by an NIH Postdoctoral Fellowship to JAC (F32AA024376), an NIH Director’s Early Independence award to WBL (1DP5OD017851-01), a Bowes Research Fellowship to WBL, and an HHMI Independent Investigator Award to MBE. LVI was supported by the Bridges to Baccalaureate program under NIH Grant R25GM095401 to Dr. Gary Firestone. We thank members of the Ludington lab, including Anjali Jain, Alison Gould, Tuzun Guvener, Benjamin Obadia, Vivian Zhang, and Carlos Zuazo, for helpful contributions to this project. We thank the laboratory of Matt Welch for use of their plate reader, April Price and the laboratory of Greg Barton for assistance with NanoStrings data collection, Henrik Jasper for an aliquot of pH3 antibody, and James Fry for help with interpretation of the internal ethanol concentration and NanoStrings results. We thank Alison Gould and Benjamin Obadia for providing helpful comments to the manuscript.

## References

Apidianakis, Y. & Rahme, L.G., 2011. Drosophila melanogaster as a model for human intestinal infection and pathology. Disease models & mechanisms, 4(1), pp.21–30.

Arrese, E.L. & Soulages, J.L., 2010. Insect fat body: energy, metabolism, and regulation. Annual review of entomology, 55(1), pp.207–225.

Blum, J.E. et al., 2013. Frequent replenishment sustains the beneficial microbiome of Drosophila melanogaster. MBio, 4(6), pp.e00860-13–e00860-13.

Broderick, N.A. & Lemaitre, B., 2012. Gut-associated microbes of Drosophila melanogaster. Gut microbes, 3(4), pp.307–321.

Broderick, N.A., Buchon, N. & Lemaitre, B., 2014. Microbiota-induced changes in drosophila melanogaster host gene expression and gut morphology. MBio, 5(3), pp.e01117–14.

Buchon, N. et al., 2009. Invasive and indigenous microbiota impact intestinal stem cell activity through multiple pathways in Drosophila. Genes & development, 23(19), pp.2333–2344.

Bull-Otterson, L. et al., 2013. Metagenomic analyses of alcohol induced pathogenic alterations in the intestinal microbiome and the effect of Lactobacillus rhamnosus GG treatment. PloS one, 8(1), p.e53028.

Chandler, J.A. et al., 2011. Bacterial communities of diverse Drosophila species: ecological context of a host-microbe model system. H. S. Malik, ed. PLoS genetics, 7(9), p.e1002272.

Chandler, J.A., Eisen, J.A. & Kopp, A., 2012. Yeast communities of diverse Drosophila species: comparison of two symbiont groups in the same hosts. Applied and environmental microbiology, 78(20), pp.7327–7336.

Chen, P. & Schnabl, B., 2014. Host-microbiome interactions in alcoholic liver disease. Gut and liver, 8(3), pp.237–241.

Clark, R.I. et al., 2015. Distinct Shifts in Microbiota Composition during Drosophila Aging Impair Intestinal Function and Drive Mortality. Cell reports, 12(10), pp.1656–1667.

Cowmeadow, R.B., Krishnan, H.R. & Atkinson, N.S., 2005. The slowpoke gene is necessary for rapid ethanol tolerance in Drosophila. Alcoholism, clinical and experimental research, 29(10), pp.1777–1786.

Devineni, A.V. & Heberlein, U., 2009. Preferential ethanol consumption in Drosophila models features of addiction. Current biology: CB, 19(24), pp.2126–2132.

Devineni, A.V. & Heberlein, U., 2013. The Evolution of Drosophila melanogaster as a Model for Alcohol Research. dx.doi.org.

Diehl, A.M., 2002. Liver disease in alcohol abusers: clinical perspective. Alcohol (Fayetteville, N.Y.), 27(1), pp.7–11.

Djawdan, M. et al., 1996. Metabolic aspects of the trade-off between fecundity and longevity in Drosophila melanogaster. Physiological Zoology, 69(5), pp.1176–1195.

Douglas, A.E., 2018. The Drosophila model for microbiome research. Lab animal, 47(6), pp.157–164.

Elya, C. et al., 2016. Stable Host Gene Expression in the Gut of Adult Drosophila melanogaster with Different Bacterial Mono-Associations. W. J. Etges, ed. PloS one, 11(11), p.e0167357.

Forsyth, C.B. et al., 2009. Lactobacillus GG treatment ameliorates alcohol-induced intestinal oxidative stress, gut leakiness, and liver injury in a rat model of alcoholic steatohepatitis. Alcohol (Fayetteville, N.Y.), 43(2), pp.163–172.

Fry, J.D., 2014. Mechanisms of naturally evolved ethanol resistance in Drosophila melanogaster. Journal of Experimental Biology, 217(Pt 22), pp.3996–4003.

Ghezzi, A., Krishnan, H.R. & Atkinson, N.S., 2014. Susceptibility to ethanol withdrawal seizures is produced by BK channel gene expression. Addiction biology, 19(3), pp.332–337.

Gibson, J.B., May, T.W. & Wilks, A.V., 1981. Genetic variation at the alcohol dehydrogenase locus in Drosophila melanogaster in relation to environmental variation: Ethanol levels in breeding sites and allozyme frequencies. Oecologia, 51(2), pp.191–198.

Gomez, C.R. et al., 2008. Innate immunity and aging. Experimental gerontology, 43(8), pp.718–728.

Gould, A.L. et al., 2018. High-dimensional microbiome interactions shape host fitness.

Guo, L. et al., 2014. PGRP-SC2 promotes gut immune homeostasis to limit commensal dysbiosis and extend lifespan. Cell, 156(1-2), pp.109–122.

Hartmann, P., Seebauer, C.T. & Schnabl, B., 2015. Alcoholic liver disease: the gut microbiome and liver cross talk. Alcoholism, clinical and experimental research, 39(5), pp.763–775.

Hockings, K.J. et al., 2015. Tools to tipple: ethanol ingestion by wild chimpanzees using leaf-sponges. Royal Society open science, 2(6), p.150150.

Ja, W.W. et al., 2007. Prandiology of Drosophila and the CAFE assay. Proceedings of the National Academy of Sciences of the United States of America, 104(20), pp.8253–8256.

Kaun, K.R. et al., 2011. A Drosophila model for alcohol reward. Nature neuroscience, 14(5), pp.612–619.

Koyle, M.L. et al., 2016. Rearing the Fruit Fly Drosophila melanogaster Under Axenic and Gnotobiotic Conditions. JoVE (Journal of Visualized Experiments), (113).

Lemaitre, B. & Hoffmann, J., 2007. The host defense of Drosophila melanogaster. Annual review of immunology, 25, pp.697–743.

Lemaitre, B. & Miguel-Aliaga, I., 2013. The digestive tract of Drosophila melanogaster. Annual review of genetics, 47(1), pp.377–404.

Lhocine, N. et al., 2008. PIMS modulates immune tolerance by negatively regulating Drosophila innate immune signaling. Cell host & microbe, 4(2), pp.147–158.

Li, H. & Jasper, H., 2016. Gastrointestinal stem cells in health and disease: from flies to humans. Disease models & mechanisms, 9(5), pp.487–499.

Logan-Garbisch, T. et al., 2015. Developmental Ethanol Exposure Leads to Dysregulation of Lipid Metabolism and Oxidative Stress in Drosophila. G3: Genes| Genomes| Genetics, 5(1), pp.49–59.

Mazeh, S. et al., 2008. The influence of ethanol on feeding in the frugivorous yellow-vented bulbul (Pycnonotus xanthopygos). Behavioural processes, 77(3), pp.369–375.

McClure, K.D., French, R.L. & Heberlein, U., 2011. A Drosophila model for fetal alcohol syndrome disorders: role for the insulin pathway. Disease models & mechanisms, 4(3), pp.335–346.

Obadia, B. et al., 2018. Diet influences host-microbiota associations in Drosophila. Proceedings of the National Academy of Sciences, 115(20), pp.E4547–E4548.

Obadia, B. et al., 2017. Probabilistic Invasion Underlies Natural Gut Microbiome Stability. Current biology: CB, 27(13), pp.1999–2006.e8.

Regan, J.C. et al., 2016. Sex difference in pathology of the ageing gut mediates the greater response of female lifespan to dietary restriction. eLife, 5, p.e10956.

Rera, M., Clark, R.I. & Walker, D.W., 2012. Intestinal barrier dysfunction links metabolic and inflammatory markers of aging to death in Drosophila. Proceedings of the National Academy of Sciences of the United States of America, 109(52), pp.21528–21533.

Ridley, E.V. et al., 2012. Impact of the resident microbiota on the nutritional phenotype of Drosophila melanogaster. F. Leulier, ed. PloS one, 7(5), p.e36765.

Ridley, E.V., Wong, A.C.-N. & Douglas, A.E., 2013. Microbe-dependent and nonspecific effects of procedures to eliminate the resident microbiota from Drosophila melanogaster. Applied and environmental microbiology, 79(10), pp.3209–3214.

Robinson, B.G. et al., 2012. Neural adaptation leads to cognitive ethanol dependence. Current biology: CB, 22(24), pp.2338–2341.

Sandhu, S. et al., 2015. An inexpensive, scalable behavioral assay for measuring ethanol sedation sensitivity and rapid tolerance in Drosophila. JoVE (Journal of Visualized Experiments), (98), pp.e52676–e52676.

Sánchez, F. et al., 2004. The possible roles of ethanol in the relationship between plants and frugivores: first experiments with egyptian fruit bats. Integrative and comparative biology, 44(4), pp.290–294.

Shaw, A.C. et al., 2010. Aging of the innate immune system. Current opinion in immunology, 22(4), pp.507–513.

Shin, S.C. et al., 2011. Drosophila microbiome modulates host developmental and metabolic homeostasis via insulin signaling. Science (New York, N.Y.), 334(6056), pp.670–674.

Skorupa, D.A. et al., 2008. Dietary composition specifies consumption, obesity, and lifespan in Drosophila melanogaster. Aging cell, 7(4), pp.478–490.

Steinfeld, H.M., 1927. Length of life of Drosophila melanogaster under aseptic conditions, Ph.D. Thesis. Department of Zoology. University of California, Berkeley.

Storelli, G. et al., 2011. Lactobacillus plantarum promotes Drosophila systemic growth by modulating hormonal signals through TOR-dependent nutrient sensing. Cell metabolism, 14(3), pp.403–414.

Tennessen, J.M. et al., 2014. Methods for studying metabolism in Drosophila. Methods (San Diego, Calif.), 68(1), pp.105–115.

Troutwine, B.R. et al., 2016. Alcohol resistance in Drosophila is modulated by the Toll innate immune pathway. Genes, brain, and behavior, 15(4), pp.382–394.

Wiens, F. et al., 2008. Chronic intake of fermented floral nectar by wild treeshrews. Proceedings of the National Academy of Sciences, 105(30), pp.10426–10431.

Wong, A.C.-N., Dobson, A.J. & Douglas, A.E., 2014. Gut microbiota dictates the metabolic response of Drosophila to diet. Journal of Experimental Biology, 217(Pt 11), pp.1894–1901.

Wood, A.M. et al., 2018. Risk thresholds for alcohol consumption: combined analysis of individual-participant data for 599 912 current drinkers in 83 prospective studies. Lancet (London, England), 391(10129), pp.1513–1523.

Xu, S. et al., 2012. The propensity for consuming ethanol in Drosophila requires rutabaga adenylyl cyclase expression within mushroom body neurons. Genes, brain, and behavior, 11(6), pp.727–739.

Zera, A.J. & Harshman, L.G., 2001. The physiology of life history trade-offs in animals. Annual review of ecology and systematics, 32, pp.95–126.

